# Characterization of potassium, sodium and their interactions effects in yeasts

**DOI:** 10.1101/2020.10.22.350355

**Authors:** Aleksandr Illarionov, Petri-Jaan Lahtvee, Rahul Kumar

**Affiliations:** Institute of Technology, University of Tartu, Estonia

**Keywords:** Microbial cell factories, *Saccharomyces cerevisiae*, *Kluyveromyces marxianus*, *Rhodotorula toruloides*, yeast, sodium, potassium, salt stress, cationic stress, osmotic stress, oxidative stress, carotenoids, food additives, image analysis, bioprocess, biotechnology

## Abstract

Biotechnology requires efficient microbial cell factories. The budding yeast *Saccharomyces cerevisiae* is an important cell factory but for a sustainable use of natural resources more diverse cellular attributes are essential. Here, we benchmarked non-conventional yeasts *Kluyveromyces marxianus* (KM) and *Rhodotorula toruloides* (RT) against the extensively characterized strains of *S. cerevisiae*, CEN.PK and W303. We developed a computational method for the characterization of cell/vacuole volumes and observed an inverse relationship between the maximal growth rate and the median cell volume that was responsive to monovalent cations. We found that the supplementation of certain K^+^ concentrations to CEN.PK cultures containing 1.0 M Na^+^ increased the specific growth rate by four-fold with a parabolic shift in the median cell/vacuole volumes. The impairment of ethanol and acetate utilization in CEN.PK, acetate in W303, at the higher K^+^/Na^+^ concentrations implied an interference in the metabolic pathways required for their consumption. In RT cultures, the supplementation of K^+^/Na^+^ induced a trade-off in glucose utilization but alleviated cellular aggregates formation where specified cationic concentrations increased the beta-carotene yield by 60% compared with the reference. Our comparative analysis of cell/vacuole volumes using exponential phase cultures showed that the median volumes decreased the most for KM and the least for RT in response to studied cations. Noteworthy for the implication in aging research using yeasts, the vacuole to cell volume ratio increased with the increase in cell volume for W303 and KM, but not for CEN.PK and RT.

**Importance:** For designing efficient bioprocesses characterization of microbial cell factories in the relevant culture environment is important. The control of cell volume in response to salt stress is crucial for the productivity of microbial cell factories. We developed an open source computational method for the analysis of optical microscopy images that allowed us to quantify changes in cell/vacuole volumes in response to common salts in yeasts. Our study provides a framework for appreciating the role of cellular/organellar volumes in response to changing physiological environment. Our analysis showed that K^+^/Na^+^ interactions could be used for improving the cellular fitness of CEN.PK and increasing the productivity of beta-carotene in *R. toruloides*, which is a commercially important antioxidant and a valuable additive in foods.

## Introduction

Microbial cell factories (MCFs) are the key drivers of biotechnology and are important for a sustainable use of resources on the planet earth. The unsustainable fossil-fuel-based chemical processes are set for replacement with the bioprocesses using MCFs (1, 2). In the industrial bioprocesses, an ideal MCF thrives under severe stress conditions such as hyperosmolarity, temperature, pH, and inhibitory effects of substrate–product concentrations in the culture environment (3–5). In particular, the budding yeast *Saccharomyces cerevisiae*, one of the most commonly used MCF, shows hypersensitivity towards salts (6). Besides poor salt tolerance, the conventional yeast *S. cerevisiae* also shows a narrow substrate–product spectrum to be applicable in a broader range of bioprocesses. Therefore, for the development of bioprocesses, it is important to identify alternative microorganisms that are more robust and benchmark those against the most common MCF. The present study is aimed at pursuing this goal by investigating tolerance and interaction of sodium and potassium cations in the non-conventional yeasts.

We used extensively characterized *S. cerevisiae* strains—CEN.PK113-7D and W303 to benchmark salt tolerance in the non-conventional yeasts—thermotolerant *Kluyveromyces marxianus* and oleaginous *Rhodotorula toruloide*s (7–13). These non-conventional yeasts show a broader substrate-product spectrum over the conventional yeast by consuming both hexose and pentose sugars that are abundantly available in the renewable natural resources and by producing specialty chemicals such as terpenoids, carotenoids and biofuels (14–16). By impacting the water activity and molecular crowding—higher concentrations of salts, nutrients, and metabolic byproducts induce ionic, oxidative, and osmotic stress in MCFs affecting cellular robustness (17–20). There are three basic mechanisms that protect cells against the adverse effects of higher solute concentrations, in particular, that of cations— the focus of the present study. Firstly, the active diffusion of cations that occurs through the molecular transporters—both at cellular and organellar levels (21–25). Secondly, the intracellular mechanisms that activate pathways, such as the high osmolarity glycerol (HOG) pathway, yielding osmolytes and antioxidants, which enhance the cellular fitness (26–28). Thirdly, the sequestration of cations within organelles that provides a permissible intracellular and organellar pH for enzymatic reactions (29–31). These molecular mechanisms are often coupled with the exchange of protons and electrochemical gradient across the biological membranes (32, 33). The difference in these underlying mechanisms enables distinct cationic responses in cells. Salt concentrations affect the biochemical and biophysical properties of cells by introducing changes in the biosynthesis and distribution of macromolecules. These molecular responses affect shape and size— both at organellar and cellular levels. The quantification of biophysical properties such as volume provides a global readout for changes introduced by varying the salt concentrations (34, 35). However, at present, a comparative description of cellular and organellar volume changes in response to salt concentrations is lacking for conventional and non-conventional yeasts but will be valuable for designing efficient MCFs in the future.

We developed a computational method for optical image analysis and applied it for characterization of sodium and potassium cationic responses in yeasts. Our study provides a quantitative description of volumetric parameters by analysing over a million datapoints in images (>820,00 cells and >290,000 vacuoles) obtained from the exponentially growing yeast cultures. Firstly, we characterized physiology and morphology of yeasts in a chemically defined culture medium. Secondly, the impact of different concentrations of potassium and sodium salts were examined individually. Thirdly, the different potassium concentrations were evaluated in a background of constant sodium concentration in the culture environment. Finally, the bioprocess application of cationic interactions was demonstrated in modulation of carotene productivity in *R. toruloides*.

## RESULTS

### Characterization of yeasts in a chemically defined culture medium

*S. cerevisiae* CEN.PK113-7D (CEN.PK), *S. cerevisiae* W303 (W303), *K. marxianus* (KM) and *R. toruloides* (RT) were cultivated in a chemically defined mineral culture medium where glucose was the main carbon source (Fig. 1A). The highest cation concentration in the reference culture medium was for K^+^ (0.025 M), followed by Mg^2+^ (0.002 M), Na^+^ (0.5×10^−5^ M), and other cations were present in the trace amounts (Table S1). We used shake flask grown cultures to determine growth and physiology differences among the studied yeasts. We found that both strains of *S. cerevisiae* grew similarly in the shake flask cultures (Fig. 1 B1-B2). KM showed the fastest growth, while RT was the slowest growing yeast (Fig. 1B1). KM attained the highest cell density among yeasts indicating the highest biomass yield (Fig. 1B2). RT formed aggregates in the reference culture medium after 7 hours of incubation, making it infeasible to reliably ascertain its cell density past that time point, and took more than six-times compared with *S. cerevisiae* to consume all the available glucose (Fig. 1B2, S1B). *S. cerevisiae* strains produced metabolic byproducts such as ethanol, acetic acid and glycerol but were able to consume those after the depletion of glucose (Fig. 1B2, S1B). Noticeably, CEN.PK produced two-times more acetic acid than W303 strain (S1B). Among the non-conventional yeasts in the reference culture medium, only observed byproduct was ethanol produced by KM (Fig. 1B2, S1B). All yeasts showed the distinct cellular morphology (Fig. 1A). We developed a computational method for the quantification of cellular morphologies based on optical microscopy images (https://github.com/a-ill/Cell-Image-Analysis-Pipeline). We performed the analysis of images (>5000 cells for each strain) obtained from the exponentially growing cultures (Fig. 1C1-2). This analysis allowed the quantification of differences in distributions of cell and vacuole volumes for the yeasts used in the study (Fig. 1C1-2). Previously, a similar analysis method showed that the variability in growth and cell division accounts for heterogeneity in the cultures (36). The median cell volume can be used to represent growth rate and cell division time. The stochasticity in growth and cell division process creates inherent noise in these parameters, but a characterization using the median cell volume distribution can suggest division and growth changes in the cultures (36). Since vacuole is critical for the control of cell volume in the presence of salts (37), we further expanded this approach by considering volume of yeast vacuole in our study (Fig. 1D1-D2). This allowed us to quantify the ratio between vacuole and cell volume that is considered important for the control of cell volume and aging in yeast (Fig. 1E) (38, 39). In our study, cell and vacuole volumes followed non-normal distribution patterns and to account for this, we independently calculated standard deviations for each half of the volume distribution (Fig. 1C1, D2). We plotted 50% of the obtained standard deviations for visualizing volume distributions for each half of data (Fig. 1C1, D2). These quantitative representations allowed us to convey precise information on the key cellular and organellar properties of yeasts and appreciate their physiological responses in our study. We found that among yeasts specific growth rate (h^−1^) differences were inversely proportional to the median cell volume, indicating smaller cell volume favored a faster growth (Fig. 1B1, C2). Our measurements were in agreement with previously reported cell volume for *S. cerevisiae* (40). One of the advantages that our image analysis method afforded was counting of the buds as potentially independent cells in the optical microscopy images which is challenging without the specific methods for counting mother and daughter cells (41). Therefore, the large number of small cells were due to potential differences in the buddying patterns among yeasts and across conditions. W303 showed the highest heterogeneity in population at the reference condition (Fig. 1C1-C2). The reason for this high cell volume heterogeneity for W303 was due to very high variability in vacuole volume (Fig. 1D2). At the reference condition, W303 and KM showed larger vacuole to cell volume ratios compared with CEN.PK and RT indicating potentially distinct vacuolar functions among yeasts in the study (Fig. 1E).

**FIG 1.**
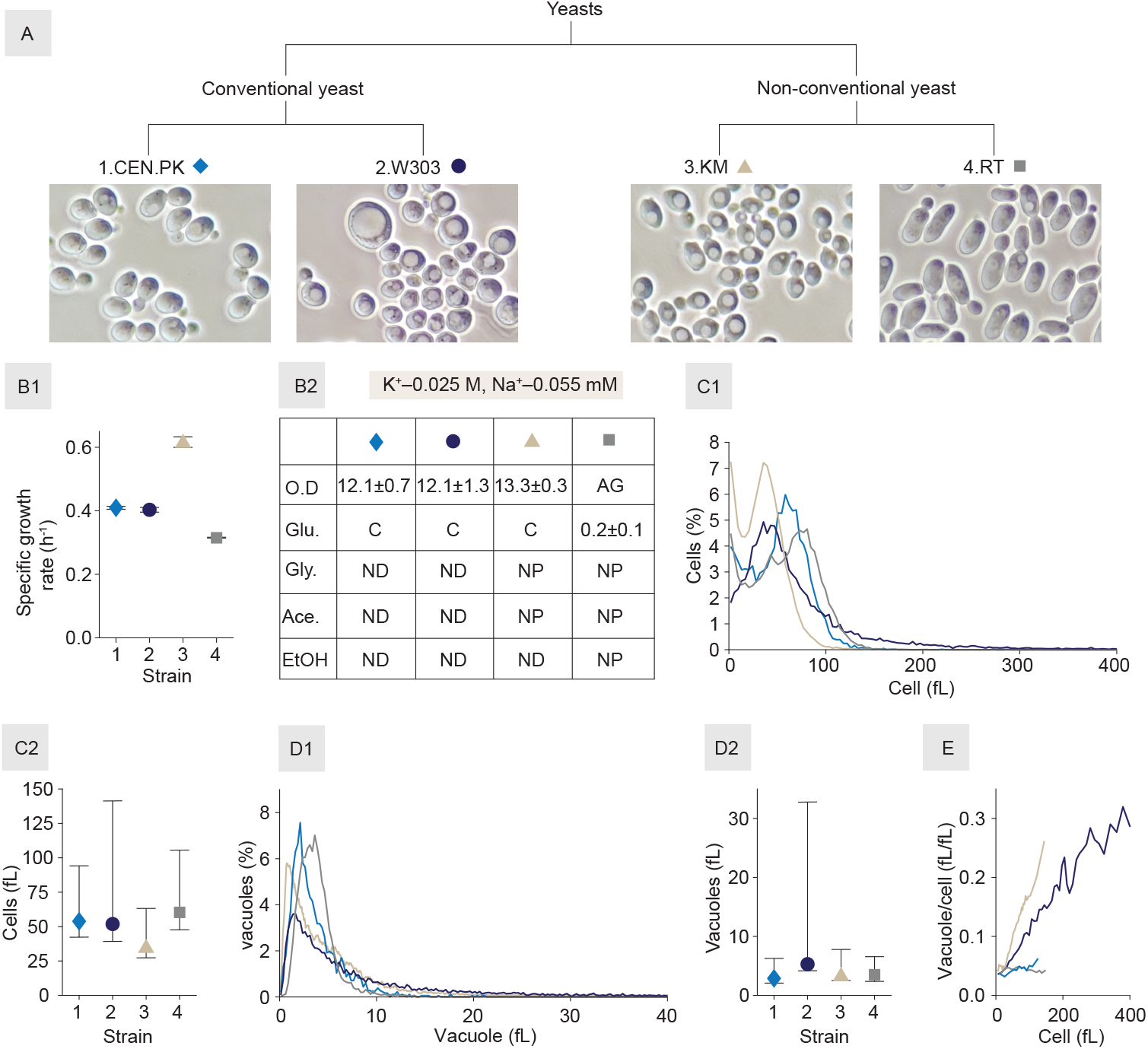
Characterization of yeasts in a chemically defined culture medium. (A) Representative morphology of exponentially growing cells; (B1) Specific growth rate (h^−1^) of yeast strains. All cultivations were performed in shake flasks. Mean values were calculated based on biological triplicate experiments; (B2) Physiological status of yeast strains at the end of culture cultivation in the reference minimal medium using shake flasks. Data represent an average of triplicate experiments and the detailed physiology profiles are available in supplemental S1B. Abbreviations: O.D= optical density at 600nm, AG= cell aggregates, Glu.= glucose (g/l), Gly.= glycerol (g/l), Ace.= acetic acid (g/), EtOH= ethanol (g/l); C= consumed, ND= produced but consumed by the end of cultivation and therefore not detected at this point, NP= Not produced; (C1) Cell volume (fL) distribution at exponential growth phase showing the percentage of cells in different volume bins. The number of cells (n) used for the quantification were as following: CEN.PK (6157), W303 (20895), KM (21534), RT (26117); (C2) Median cell volume (fL, represented by a symbol) of exponentially growing cells. Bars represent standard deviations (SD) and were calculated separately for each half of non-normal cell volume distribution as represented in C1. Both halves of SD values are plotted at 50% of the actual values for a representative visualization; (D1) Vacuole volume (fL) distribution at the exponential growth phase showing the percentage of vacuoles in different volume bins. The number of vacuoles (n) used for the quantification were as following: CEN.PK (1825), W303 (14634), KM (15412), RT (11777); (D2) Median vacuole volume (fL, represented by a symbol) of exponentially growing cells. Bars represent standard deviations as described in C2; (E) Median vacuole to cell volume ratio at exponential growth phase. The number of vacuoles and cells used to calculate this ratio is the same as in D1.

### Effects of potassium salt supplementation in yeasts

After the characterization of yeasts in the reference culture medium, we used a microplate reader set up for the screening and identification of the viable K^+^ concentration range in the study. In the screening process, K^+^ concentration was varied from the reference level to 2.0 M by using KCl (Fig. 2). We found that yeasts were viable up to 1.5 M K^+^ concentration but were unable to survive at the next tested K^+^ concentration i.e., 2.0 M, except for RT (Fig. 3A). KM showed the highest sensitivity to increase in K^+^ concentration by reducing specific growth rates by 25% at 1.0 M and 58% at 1.5 M, respectively (Fig. 2A). However, since the reference specific growth rate of KM was the highest among the studied yeast it nevertheless maintained the highest specific growth rate at viable K^+^ concentrations (Fig. 1B1, 2A). KM was followed by CEN.PK for the sensitivity towards K^+^ as it showed 11% and 60% reduction in specific growth rate in comparison with the reference at 1.0 M and 1.5 M concentration, respectively (Fig. 1B1, 2A). RT was only viable strain at 2.0 M K^+^ concentration where it grew at 54% reduced specific growth rate compared with the reference cultivation condition, indicating a higher tolerance for K^+^ cations (Fig. 2A). Thereafter, we used shake flask cultivations to determine physiological responses in the presence of 1.0 M K^+^ concentration in the cultures where all the strains maintained 75% or more of specific growth rate compared with the reference condition (Fig. 2B, S1B). *S. cerevisiae* strains responded differently to increase in K^+^ concentration. Although the consumption of glucose in CEN.PK was not affected, it failed to consume the metabolic byproducts accumulated during growth on glucose. Interestingly, CEN.PK converted the produced ethanol to acetic acid, but the latter was not further metabolized (Fig. 2B, S1B). However, W303 consumed the metabolic byproducts, except glycerol which was consumed only partially during the cultivation duration (Fig. 2B, S1B). Unlike CEN.PK, W303 strain showed no impairment upon the increased K^+^ concentration (S1B). As a consequence of these metabolic disparities CEN.PK and W303 showed 65% and 37% lower cells density, respectively, compared with the references cultures where both strains showed similar cell densities (Figure 2B, S1B). Again, KM showed the maximum cell density among the strains, despite the decrease of 33% compared with the reference condition. It slowly but completely exhausted glucose with glycerol being the main byproduct which was also depleted by the end of cultivation (Fig. 2B, S1B). RT showed a remarkable trade-off between the consumption of glucose and formation of cellular aggregates in the culture (Fig. 2B, S1B). The addition of 1.0 M KCl eliminated aggregates from RT culture but led to a slower and only a partial consumption of glucose (Fig. 2B, S1B). It also showed a small accumulation of glycerol in the culture broth but no production of acetic acid or ethanol (Fig. 2B, S1B). For the quantification of cellular and vacuolar volumes, we investigated a viable and less detrimental K^+^ concentration range (0.025–1.0 M) for growth by using shake flask cultivations and collected samples for the image analysis at exponential phase (Fig. 2C, S2A, S2B). For CEN.PK, heterogeneity in both median cellular and vacuolar volumes remained nearly constant, but the median volumes showed a significant change at higher K^+^ concentration (Fig. 2C, S2A, S2B). This was in contrast to the response of W303, the most heterogenous of populations in the study, that showed a significant increase in both median cellular and vacuolar volumes at the increased K^+^ concentrations (Fig. 2C, S2A, S2B). The vacuole to cell volume ratio slope did not change significantly for both *S. cerevisiae* strains. KM showed an exponential decrease in median cell volume but a U–shaped pattern for median vacuolar volume—both of which changed significantly with the increase in K^+^ concentration. For KM, the overall population-level heterogeneity for both cells and vacuoles initially appeared unchanged but increased at the highest concentration (Fig. 2C, S2A, S2B). For KM, vacuolar volume did not follow the pattern of decrease in cell volume with increase in K^+^ concentration. Initially, KM vacuole volume decreased with the increase in K^+^ concentration, but at 1.0 M K^+^ it partially reverted towards the reference (Fig. 2C, S2A, S2B). The increase in the vacuole to cell volume ratio slope with the increase in cell volume, which was present at the reference condition disappeared with addition of 0.3-0.6 M of K^+^ concentrations. At 1.0 M K^+^ the increase in slope reappeared, but with a smaller angle compared with the reference. This increase in the slope appeared be connected to the increase in vacuole volume at 1.0M K^+^ concentration (S2B, Fig. 1E, Fig. 2C). For RT both cellular and vacuolar median volumes significantly increased in the similar direction and it showed the least imparted heterogeneity due to the increase in K^+^ concentration among the studied strains (Fig. 2C, S2A, S2B). The vacuole to cell volume ratio slope remained unaffected by K^+^ supplementation in RT cultures (S2B).

**FIG 2.**
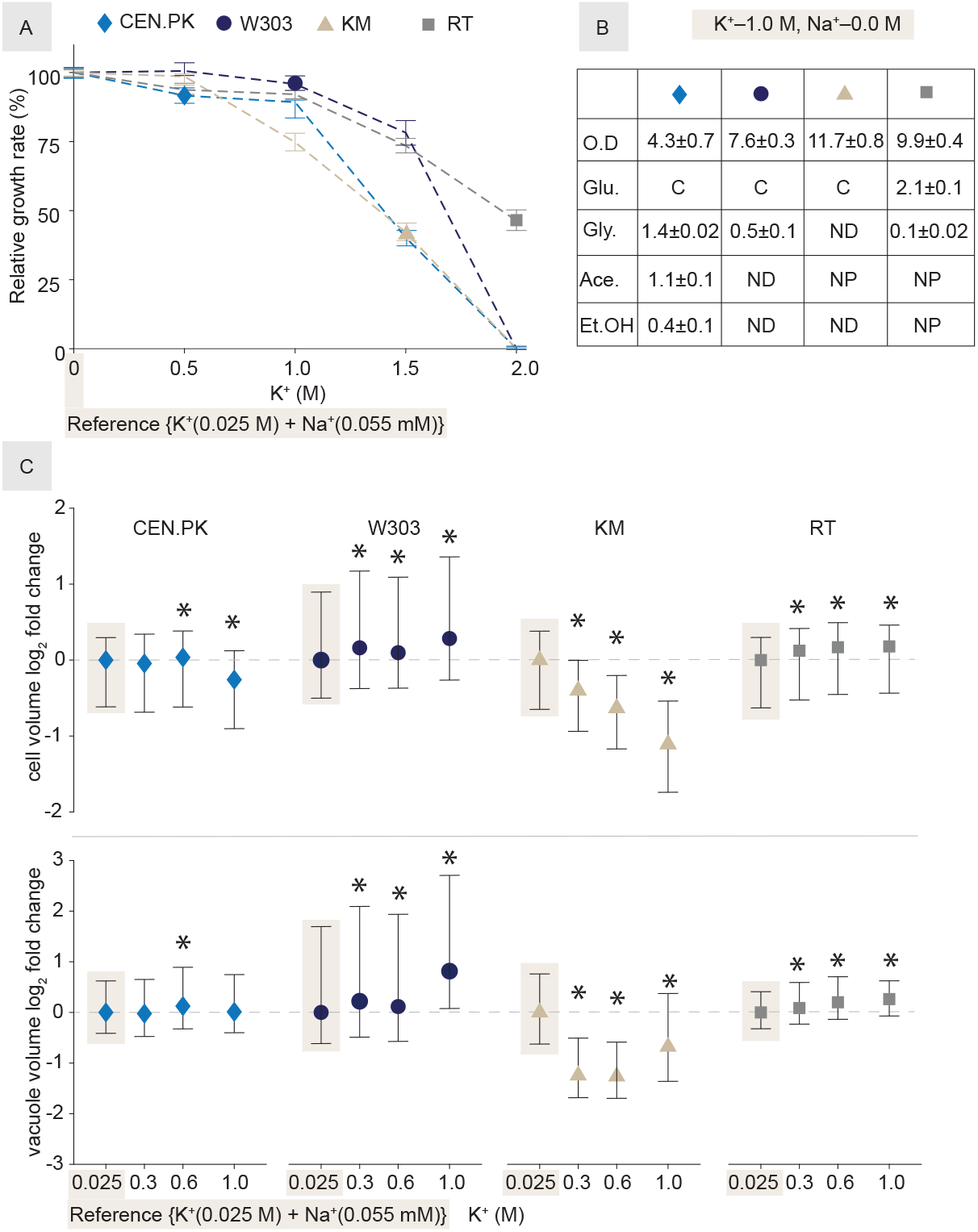
Response of yeast strains to the supplementation of K^+^ in a chemically defined culture medium. (A) Relative specific growth rate, using microplate reader, based on independent quadruplicate experiments. Specific growth rates in the reference culture medium (K^+^–0.025 M, Na^+^–0.0 M,) were considered at 100 percent for each yeast. Error bars indicate SD; (B) Physiological status of yeast strains at the end of cultivation in the cation supplemented reference minimal medium using shake flasks. Data represent average of triplicate experiments and detailed physiology profiles are available in supplemental S1B and corresponding cellular morphology in S2B. Abbreviations are the same as used in Figure 1; (C) Relative cell (n>6000) and vacuole (n>1800) volume distributions of exponential phase cells. The first data point (indicated by a shadowed background) is the reference (refers to Figure 1). Log_2_ fold changes (log_2_FC= median values, indicated by symbols) are plotted relative to the reference. Bars representing log_2_FC(SD) are calculated separately for each half of non-normal cell and vacuole volume distributions, respectively. A log_2_FC is obtained by dividing SD of experimental condition with the reference value for each half of non-normal distribution, separately. Both halves of the SD values are plotted at 50% of the actual values for a representative data visualization. Supplemental 2B shows percentage bins of cell and vacuole distributions in response to the increased K^+^ concentration in the culture medium. Asterisks (*) indicate statistical significance (p–value <0.001) based on the Kruskal-Wallis test, which is a modified version of one-way analysis of variance (ANOVA) and applicable to non-normal distributions.

**FIG 3.**
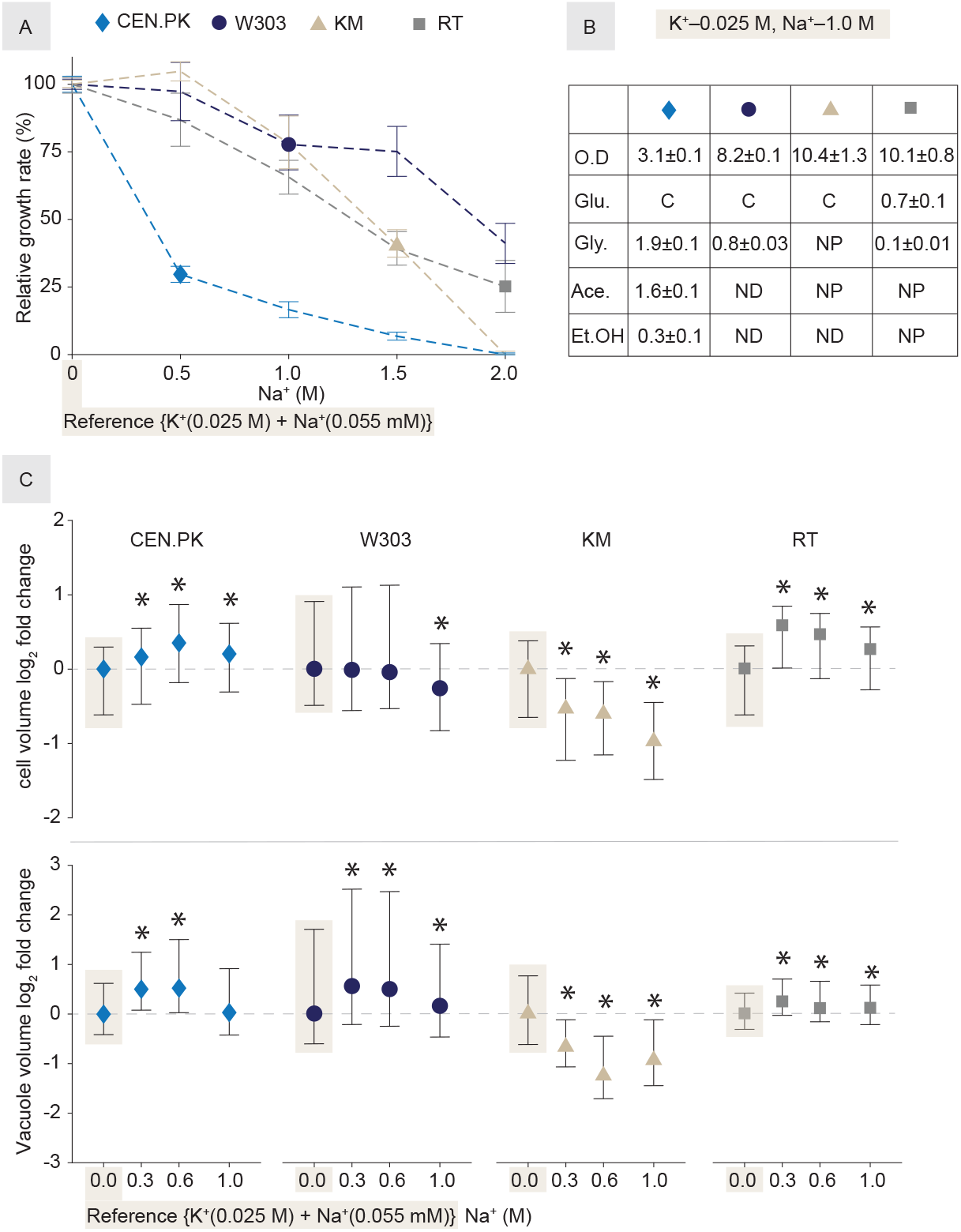
Response of yeast strains to the supplementation of Na^+^ in a chemically defined culture medium. (A) Relative specific growth rate, using microplate reader, based on independent quadruplicate experiments. Growth rates in the reference culture medium (Na^+^–0.0 M, K^+^– 0.025 M) are considered at 100 percent for each yeast and changes in growth rates (mean values) are plotted as relative to the reference. Error bars indicate SD; (B) Physiological status of yeast strains at the end of cultivation in Na^+^ supplemented minimal medium using shake flasks. Data are average of triplicate experiments and detailed physiology profiles are available in the supplemental S1B and corresponding cellular morphology in S3A. Abbreviations are the same as used in Figure 1; (C) Relative cell (n>6000) and vacuole (n>1500) volume distributions of exponential phase cells. The first data point (indicated by a shadowed background) is the reference (refers to Figure 1). Log_2_ fold changes (log_2_FC= median values, indicated by symbols) are plotted relative to the reference. Bars representing log_2_FC(SD) are calculated separately for each half of non-normal cell and vacuole volume distributions, respectively. A log_2_FC is obtained by dividing SD of experimental condition with the reference value for each half of non-normal distribution, separately. Both halves of the SD values are plotted at 50% of the actual values for a representative data visualization. The supplemental 3B shows percentage bins of cell and vacuole distributions in response to the supplementation of Na^+^ in the minimal culture medium. Asterisks (*) indicate statistical significance (p–value <0.001) based on the Kruskal-Wallis test which is a modified version of one-way analysis of variance (ANOVA) and applicable to non-normal distributions.

### Effects of sodium salt supplementation in yeasts

Similar to the investigation of K^+^ supplementation, firstly, we determined the limit of viable Na^+^ concentration using a microplate reader-based set-up, while maintaining K^+^ concentration at the reference level (Fig. 3A). We observed a drastically different response to the increased Na^+^ concentration in *S. cerevisiae* strains (Fig. 3A). The specific growth rate of CEN.PK reduced by 75% at 0.5 M Na^+^ concentration whereas W303 grew at 59% reduced specific growth rate even at 2.0 M Na^+^ concentration (Fig. 3A). This was in contrast to 2.0 M K^+^ concentration where both *S. cerevisiae* strains, along with KM, were non-viable (Fig. 2A). In the presence of 1.0 M K^+^ all strains in the study were able to grow within 75% of the specific growth rate, but in the presence of 1.0 M Na^+^ only KM and W303 maintained growth in this range while RT and CEN.PK grew at 66% and 17% of the reference specific growth rates, respectively (Fig. 2A, 3A). In the comparison of specific growth rates, KM appeared similarly sensitive towards both K^+^ and Na^+^ cations, while RT showed more sensitivity for Na^+^ compared with K^+^ cations (Fig. 2A, 3A). Secondly, in a comparable set-up to K^+^ experiments, we used shake flask cultivations supplemented with 1.0 M Na^+^ concentration to determine physiological responses of the strains in the study (Fig. 3B, S1B). *S. cerevisiae* strains responded differently to the increase in Na^+^ compared with K^+^ cation concentration (Fig. 3A, 2A, S1B). In the presence of 1.0 M Na^+^ in shake flasks, CEN.PK and W303 showed 25% and 68% of cell density, respectively, compared with the reference cultivation conditions, which were 10% less for CEN.PK but about 5% more for W303 compared with K^+^ cultivation in a comparable condition (Fig. 1B2, 2B, 3B, S1B). CEN.PK showed even more residual metabolic byproducts (33% ethanol, 35% glycerol and 45% acetic acid) in the presence of 1.0 M Na^+^ vs. 1.0 M K^+^ but similarly failed to consume those (Fig. 3B, S1B). In contrast to CEN.PK, W303 consumed both ethanol and acetic acid but only showed a partial consumption of glycerol at the end of cultivation (Fig. 3B, S1B). W303 showed nearly 60% increase in the residual glycerol in the presence of 1.0 M Na^+^ compared with 1.0 M K^+^ concentration at the end of cultivation which was nevertheless less compared with CEN.PK (Fig. 3A). Overall these metabolic disparities resulted in different biomass formation for the strains of *S. cerevisiae* (Figure 3B, S1B). In comparison to *S. cerevisiae* strains, KM showed no accumulation of metabolic byproducts at the end of cultivation (Fig. 3B, S1B). KM showed the maximum cell density among the cultures at the end of cultivation and 10% difference between cells densities in Na^+^ and K^+^ supplemented cultures (Fig. 3B, S1B). KM did not show any glycerol production in the presence of Na^+^ cations which was noticed for K^+^ cation, but ethanol metabolism appeared to be similarly disposed for both cations (Fig. 3B, S1B). RT showed a similar physiological response in the presence of Na^+^ as it showed for K^+^, except for the consumption of glucose that it consumed more in response to the increase in Na^+^ than K^+^ but with a similar accumulation of glycerol (Fig. 3B, S1B). Similar to KCl, the addition of NaCl also eliminated aggregates formation in RT cultures (Fig. 3B, S1B). For the quantification of cellular and vacuolar volumes, we investigated a viable Na^+^ concentration range (0.5×10^−5^–1.0 M) by using shake flasks and performed the image analysis on samples collected at exponential phase (Fig. 3C, S3A, S3B). For CEN.PK, both median cellular and vacuolar volumes showed a parabolic pattern though population heterogeneity remained nearly constant with the increase in Na^+^ concentration (Fig. 3C, S3A, S32B). In contrast to CEN.PK, W303 responded opposingly both at 1.0 M Na^+^ and 1.0 M K^+^ concentration with vacuole volume being significantly important contributor towards cell volume and overall heterogeneity in population (Fig. 3C, 2C). The vacuole to cell volume ratio slope did not significantly change for CEN.PK, but for W303 the increase in the slope of the vacuole to cell volume ratio increased with the increase in Na^+^ concentration (S3B). Similar to K^+^ cultivations, KM showed a significant decrease in both median cellular and vacuolar volumes with the increase in Na^+^ concentration with a relatively steady proportion of heterogeneity in population (Fig. 3C, S3A, S3B). The supplementation of Na^+^ cations affected KM similar to K^+^, where the increase in the vacuole to cell volume ratio with the increase in cell volume was not be observed for 0.3 and 0.6 M Na^+^ concentrations, but it reappeared at 1.0 M concentration with a smaller slope angle and was likely connected to the increase in vacuole volume (S3B, Fig. 1E, Fig. 3C). In response to the supplementation of Na^+^, RT also showed a parabolic pattern similar to CEN.PK for median cellular volume and significantly increased median vacuolar volume but maintained the least heterogeneity among yeasts in the study (Fig. 3C, S3A, S3B). For RT, though vacuolar distribution patterns were similar both in the presence of Na^+^ and K^+^ cations, but cell volume changes were much more apparent in response to Na^+^ than K^+^ cation (Fig. 3C, 2C). The vacuole to cell volume ratio remained unaffected by K^+^ supplementation in RT cultures (S3B).

### Modulation of sodium salt effects by supplementing potassium salt in yeasts

The distinct responses to the supplementation of K^+^ and Na^+^ cations prompted us to investigate the response of yeasts to their interactions in our study. In this instance, we used shake flask cultures for characterization of growth, physiology and morphology, and used a viable range of concentrations that were determined in the single cation supplementation experiments (Fig. 4A-C, S4A, S4B, S4C, S1B). Here, we varied K^+^ concentrations in the background of 1.0 M Na^+^ concentration to probe K^+^–Na^+^ interactions in yeasts (Fig. 4A-C). Interestingly, we found a four-fold increase in specific growth rate of CEN.PK as a result of K^+^–Na^+^ interaction (Fig. 4A). Remarkably, this four-fold increase originated by two different cellular and vacuolar volumetric patterns wherein volumes initially increased with the increase in K^+^ but reverted towards the reference (1.0 M Na^+^) levels at 0.6 M K^+^ supplementation (Fig. 4A, 4C). This was in contrast to the reference condition (without additional cation supplementation) where we observed an inverse relationship between cell volume and growth rate (Fig. 1B1, 1C2). However, this relationship was observed in W303 for K^+^-Na^+^ interactions where cellular and vacuolar volumetric increases correlated with the decrease in specific growth rate (Fig. 4A, 4C). The vacuole to cell volume ratio slope did not significantly change for *S. cerevisiae* strains (S4C). For KM, initial increase in cell and vacuole volumes correlated with the decrease in specific growth rate but at 0.4 M K^+^ supplementation significant decrease in the volumetric parameters did not correlate with the increase in growth rate (Fig. 4A, 4C). The vacuole to cell volume ratio slope increased with the increase in cell volume and persisted with the 0.1 and 0.2 M K^+^ supplementation, but a further addition of K^+^ did induce the increase in slope and it correlated with a significant decrease in vacuole volume (Fig. 4C, S4C). We found that though for single cations KM tolerated up to 1.0 M concentrations, but in K^+^-Na^+^ interaction experiments it showed a remarkable extension of the lag phase at viable K^+^ concentration and the supplementation of 0.6 M K^+^ concentration was lethal for it (Fig. 4A, S4A, S4B). In the K^+^-Na^+^ interaction experiments, RT showed the most robust growth where supplementation up to 0.4 M K^+^ did not decrease growth rate and even at 0.6 M K^+^ concentration it maintained 82% of specific growth rate observed for the 1.0 M Na^+^ supplemented cultures. Interestingly, this maintenance of specific growth rate was accompanied by a significant decrease in the measured volumetric parameters for RT (Fig. 4A, 4C, S4A, S4B). The supplementation of K^+^ in 1.0 M Na^+^ background reduced cell density in yeasts compared with single cation concentrations (Fig. 4B). CEN.PK failed not only to consume ethanol but, unlike in the presence of single cations, was also impaired in its ability to convert it to acetic acid (Fig. 4B, S1B). Additionally, the supplementation of K^+^ led to more than one and half-times higher glycerol production and none of it was consumed by CEN.PK indicating a complete inability to consume respiratory carbon sources in the presence of prohibitive concentrations of K^+^–Na^+^ cations in the cultures (Fig. 4B, S1B). However, W303 showed the ability to consume ethanol which led to two-times more glycerol and acetic acid accumulation compared with single cationic stress and neither of the latter carbon sources were consumed (Fig. 4B, S1B). In comparison to *S. cerevisiae* strains, KM produced and consumed glycerol and acetic acid while completely exhausting available glucose under the increased K^+^–Na^+^ concentrations where it was viable but reduced cell density by 43% compared with single cation supplementations (Fig. 4B, S1B). In similar cultivation conditions, RT showed only slight decrease in cell density compared with single cation supplementations but accumulated three-times more glycerol while showing only partial glucose consumption similar to K^+^ only supplementation indicating the latter cation adversely impacted glucose metabolism in RT (Fig. 4B, S1B). Similar to single cationic stress conditions, the vacuole to cell volume ratio remained unaffected in RT cultures (S4C).

**FIG 4.**
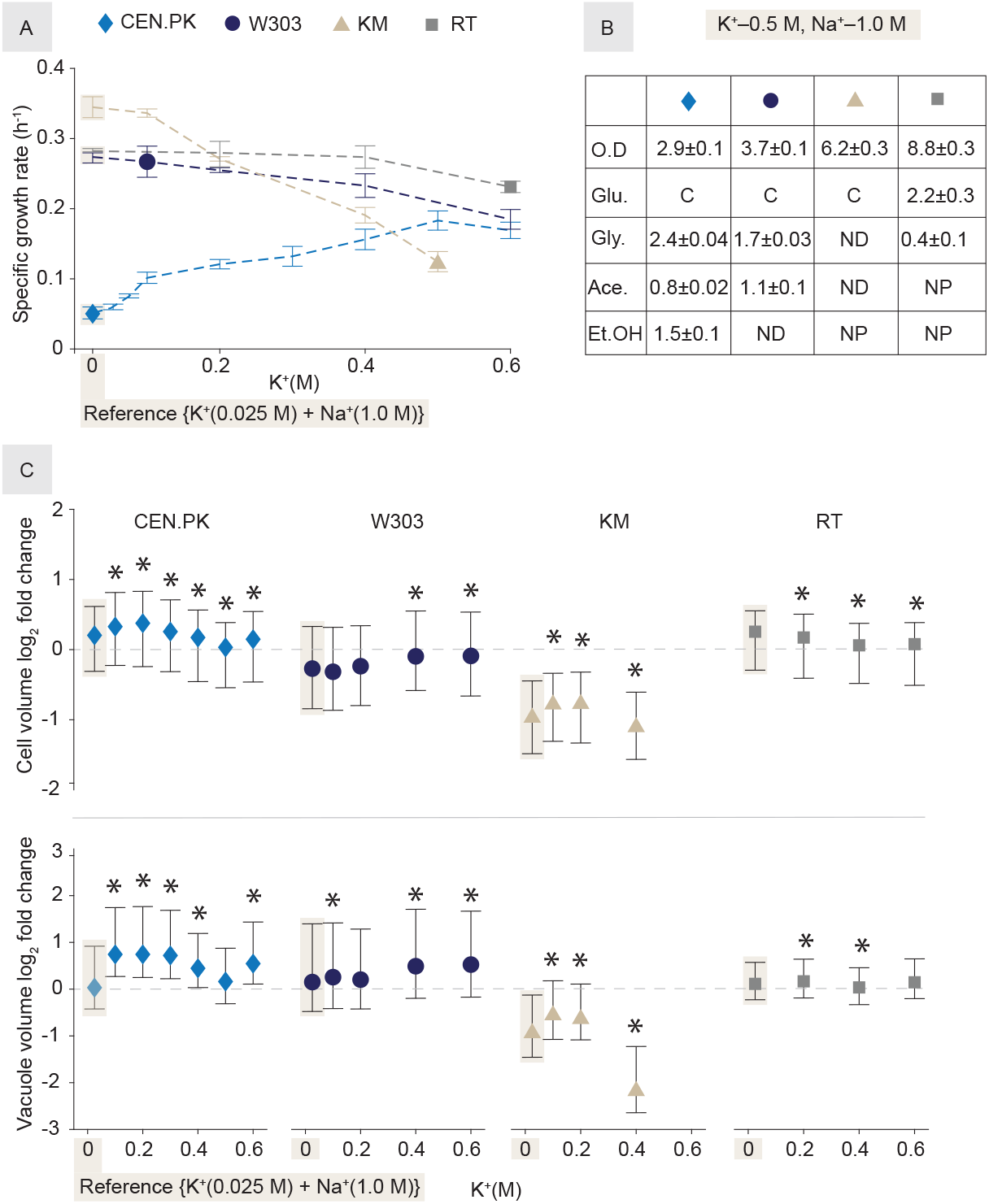
Response of yeast strains to K^+^-Na^+^ supplementation in a chemically defined culture medium. All experiments were performed in the background of 1.0 M Na^+^ while varying K^+^ concentrations. (A) Specific growth rate using shake flasks, based on independent triplicate (for CEN.PK and W303) and duplicate (KM and RT) experiments. Error bars indicate SD. Supplementary S4A shows changes in the lag phase and the final cells densities in response to dual cationic stress; (B) Physiological status of yeast strains at the end of cultivation in the K^+^-Na^+^ supplemented minimal medium using shake flasks. Data are average of triplicate experiments and detailed physiology profiles are available in supplemental S1B and corresponding cellular morphology in S4B. Abbreviations are the same as used in Figure 1; (C) Relative cell (n>7000) and vacuole (n>900) volume distributions at exponential phase. The first data point (indicated by a shadowed background) is the reference (refers to Fig. 3— K^+^– 0.025 M, Na^+^–1.0 M,). Log_2_ fold changes (log_2_FC= median values, indicated by symbols) are plotted relative to the reference. Bars representing log_2_FC(SD) are calculated separately for each half of non-normal cell and vacuole volume distributions, respectively. A log_2_FC is obtained by dividing SD of experimental condition with the reference value for each half of non-normal distribution, separately. Both halves of the SD values are plotted at 50% of actual values for a representative data visualization Supplemental 4C shows percentage bins of cell and vacuole distributions in response to K^+^-Na^+^ supplementation in the minimal medium. Asterisks (*) indicate statistical significance (p–value <0.001) based on the Kruskal-Wallis test which is a modified version of one-way analysis of variance (ANOVA) and applicable to non-normal distributions.

### K^+^ modulates carotene production in Na^+^ supplemented *R. toruloides* cultures

Toxicity in the culture environment impacts titers and productivity in bioprocesses (42). The potassium anti-porter modulated K^+^/H^+^ electrochemical gradient across membrane is proposed to be a general mechanism for enhancing tolerance to multiple alcohols when using *S. cerevisiae* as microbial cell factory (42). Here, we were interested in identifying whether K^+^ and Na^+^ symporters could also be used similarly for improving the productivity in bioprocesses. The availability of these cationic transporters differed in the studied yeasts (Fig. 5A, S5A-B). In our analysis, we found that RT lacked some of the organellar transporters, namely Mdm38, Vnx1, and Vhc1 that were present in the other yeasts (Fig. 5A, S5A-B) (9, 12, 43, 44). As RT is a native producer of carotenes and previously showed the increased production of carotenoids in the presence of osmotic stress (16, 28), we investigated whether potassium and sodium salts induced osmotic-stress can modulate the carotene titers. We found that the supplementation of 1.0 M K^+^ did not significantly improve carotene productivity (mg beta-carotene/g glucose) but the supplemental 1.0 Na^+^ concentration induced a significant (p_value_ <0.05) improvement in the productivity of beta-carotene in glucose grown cultures compared with the reference (Fig. 5B). Interestingly, the addition of 0.1 M K^+^ to 1.0 M Na^+^ containing cultures further enhanced the productivity of beta-carotene (p_value_ <0.05) by 60% compared with the reference (Fig. 5B). However, a subsequent increase in K^+^ concentration reduced the productivity of beta-carotene, indicating a limit to the cationic stress tolerance in RT (Fig. 5B). In glucose grown RT cultures, we consistently observed cellular aggregates formation in the reference culture medium but not in the presence of higher cationic concentrations. Therefore, we tested a range of K^+^ and Na^+^ cationic concentrations to identify the minimal possible concentration that disallows formation of cellular aggregates in the RT cultures. We identified that the supplementation of either 0.2 M K^+^ or 0.2 M Na^+^ to the reference cultures was sufficient to disallow formation of cellular aggregates without impacting the growth. The cell density presented for the reference was obtained in the presence of 0.2 M K^+^ concentration (Fig. 5B).

**FIG 5.**
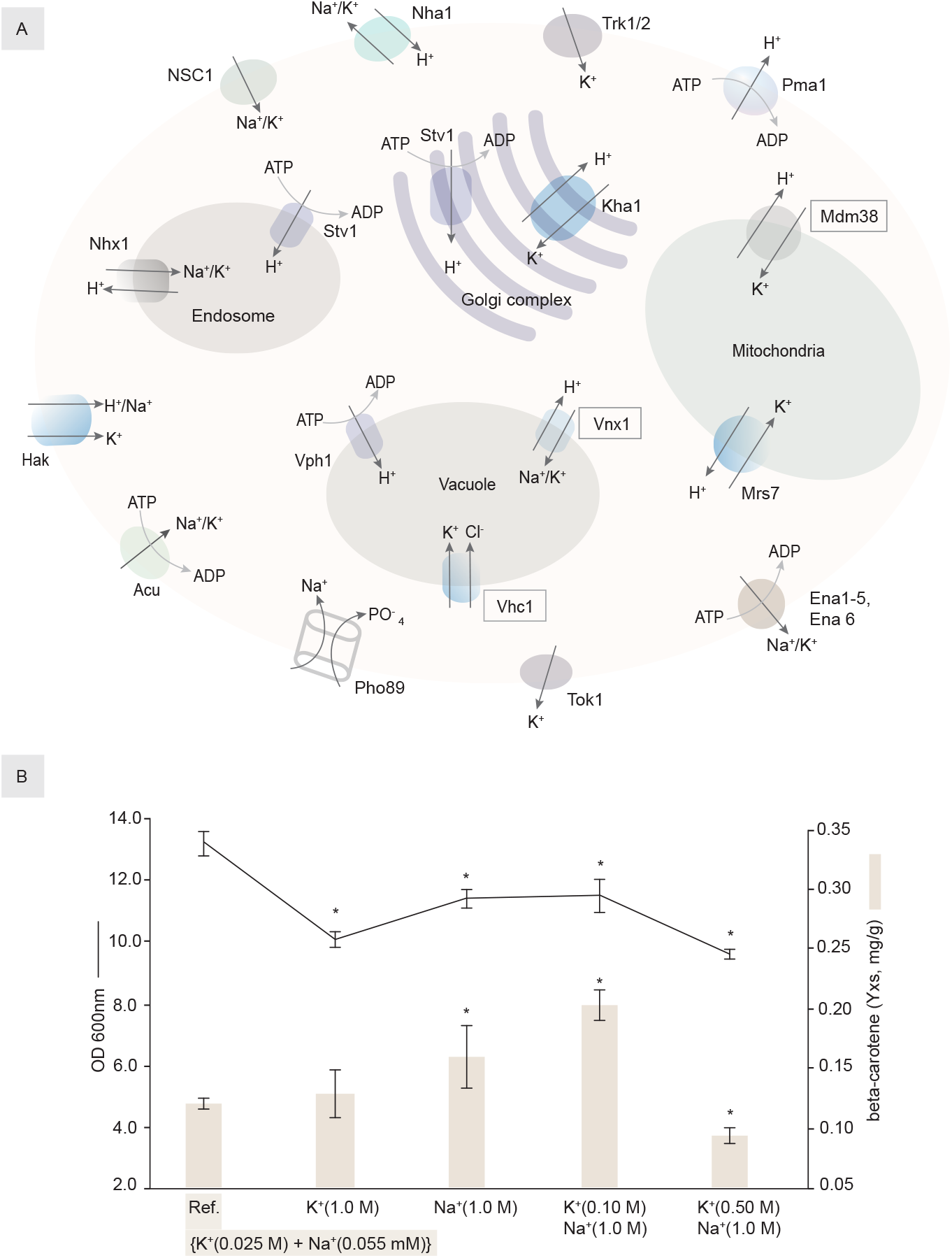
Evaluation of K^+^/Na^+^ supplementation impact on the production of beta-carotene in *R. toruloides*. (A) Cellular and organellar cationic transporters in yeasts. The absence of cationic transporters, based on the currently available sequence comparisons, in *R. toruloides* is indicated by placing the label inside a rectangle. (B) Cell density of glucose grown *R. toruloides* cultures and the beta-carotene production yield on glucose (Yxs, mg/g) in response to the supplementation of K^+^/Na^+^ cations. Data presented are from independent triplicate experiments and beta-carotene measurements are performed using the stationary phase cultures where all glucose (10 g/l) was consumed. Error bars indicate SD and significance (p_value_ <0.05) is indicated by a symbol (*).

## Discussion

Our study investigates monovalent cations (K^+^, Na^+^ and K^+^–Na^+^ interactions) impact in conventional and non-conventional yeasts and describes their physiological and morphological features. It indicates the potential application of cations in the modulation of cellular fitness and productivity in bioprocesses. We showed that K^+^ modulates toxic effects of Na^+^ cations that can be applied to improve the salt tolerance in CEN.PK and the yield of beta-carotene in RT. The supplementation of K^+^ improves the tolerance for multiple stresses, including for alcohol in yeasts (42, 45, 46). For efficient functioning of cellular processes, such as protein synthesis, high intracellular ratio K^+^/Na^+^ is essential in eukaryal organisms (19, 47). However, in the industrial environments, Na^+^ is often present at a higher concentration where it is toxic to yeast because of its ability to readily replace K^+^ in the essential cellular processes (3, 19). In particular, some of the differences towards stress tolerance are due to the specific efflux and influx transporters among yeasts (S5A-B) (25). The hyper-sensitivity of CEN.PK to Na^+^ but relative insensitivity to the same cation concentration in W303 is attributable to the presence of an atypical plasma membrane ATPase related (*PMR2*) locus, renamed as *ENA* – *exitus natru* or sodium exit (6, 21). The four-fold increase in specific growth rate of CEN.PK by the supplementation of K^+^ in the presence 1.0 M Na^+^ concentration is likely due to the differences in cation transporters among *S. cerevisiae* strains (S5A-B) (6). The role of monovalent cations in *S. cerevisiae* XT300.3A indicates requirement of a certain K^+^ concentration (0.1-0.5 mM) for growth and suggests a modulation of growth upon the supplementation of Na^+^ (0.0-150 mM) or Rb^+^ (0.0-30.0 mM) cations (48). The range of investigated cations, indicated in the brackets, are much lower than the present study but our results indicate to a similar conclusion concerning growth of CEN.PK. The reason for such growth modulation appears to be the role of intracellular K^+^ in counteracting the toxicity caused by Na^+^ at a higher concentration (49). Though the most transporter proteins consist of the same amino acid sequences between W303 and CEN.PK strains, K^+^–importers Trk1p and Trk2p show distinct peptide compositions (S5A-B). An atomic-scale model of these K^+^–importers using *S. cerevisiae* BY4741, the parental strain of CEN.PK and W303, shows that the amino acid specificity is not only important for the function but also essential for the correct positioning in membrane (50). Interestingly among the non-conventional yeasts, KM shows only a single copy of the Trk family transporter, while RT possesses both Trk1p and Trk2p (S5A-B). KM and RT also possess additional cationic transporters namely the Hak family transporters compared with *S. cerevisiae* (S5A-B). RT shows even more distinctiveness concerning the cationic transporters in having the Acu family and additionally consisting twice the number of main plasma membrane K^+^ and Na^+^ exporters, namely Tok1p and Nha1p as compared to other yeasts (S5A-B). Moreover, since organellar volume is important for cellular heterogeneity it is noteworthy that RT shows the least amount of population heterogeneity potentially because it lacks vacuole transporters such Vhc1p and Vnx1p, as well mitochondrial K^+^ transporter Mdm38p, but instead, consist of an additional copy of K^+^ transporter in Golgi complex (S5A-B). These cellular and organellar cationic attributes may contribute towards a relative population homogeneity and robustness in RT, making it an attractive candidate for deploying as MCF. Further, yeasts also improve tolerance for multiple stresses by increasing the biosynthesis of lipids which are important for membrane integrity and might be an important factor for the apparent robustness exhibited by RT in our study since it is a known oleaginous yeast that produces more lipids in the osmotic and oxidative stress environments (16, 51). The increase in the yield of antioxidant beta-carotene by 60% upon the supplementation of certain K^+^-Na^+^ cations appears to be induced in response to cationic stress (Fig. 5B). Interestingly, the increase in either K^+^ or Na^+^ concentration eliminates formation of cellular aggregates in RT, which it uses in the native environment to adhere to plant surfaces (52). This aggregate formation is disrupted by the presence of alkali-soluble materials such as mannose residues on the cellular membrane and the role of either K^+^/Na^+^ appears relevant to this disruption in our study (52). Another factor that contributes to the osmotic tolerance is the biosynthesis of osmoprotectant such as glycerol (26). We observed differences in the production and consumption of glycerol in *S. cerevisiae* strains and RT, compared with KM. KM produced much less glycerol compared with other strains in the study and consumed it completely unlike other strains in the presence of higher salt concentrations. KM harbors the evolutionarily conserved HOG pathway and produces much higher levels of glycerol when lactose is used as a carbon source as compared with glucose (27). Although we lack a direct comparison, these observations suggest that the HOG pathway regulation and glycerol metabolism in KM may potentially be carbon source dependent which will be consistent with the dependency of osmostress response on carbon sources as demonstrated for W303 by growing it on glucose and ethanol in the presence of NaCl (53). Also concerning the utilization of carbon sources, *S. cerevisiae* strains produce the metabolic byproducts such as ethanol and acetic acid while consuming glucose, but during the diauxic shift these byproducts tend to be consumed (54). However, we observed that the higher K^+^/Na^+^ concentrations, both individually and together, impinged on ethanol and acetic acid utilization in CEN.PK, and when both of the cationic concentrations were increased, W303 also failed to fully consume acetic acid. In aerobic yeast cultures, ethanol and acetic acid are utilized via gluconeogenesis and together with glycerol, which is utilized via glycolytic pathway, require respiratory metabolism for their consumption that is affected by osmotic stress (3). It is plausible that K^+^/Na^+^ cations not only have an effect on ion and solute exchanges across the mitochondrial membrane but also on enzyme activities involved in the utilization of energy sources (47). The metabolic differences between CEN.PK and W303 are interesting for understanding K^+^/Na^+^ interaction dynamics and mechanisms involved in the osmostress response in *S. cerevisiae*. Finally, we observed an inverse relationship between cell volume and maximum growth rate among yeasts which was responsive to cationic interventions for a given strain used in the study. The relationship in growth rate and cell volume is likely due to differences in the genomic attributes among yeasts (as noted under strains and culture conditions section). We observed a large number of the smaller cells in yeast populations which were likely related to the impact on formation of buds that besides being important in cell division are also relevant to the cytosolic pH control as nascent daughter cells lack the plasma membrane proton ATPase (Pma1) (Fig. 5A) (41, 55). As yeasts are often used in the aging research, our findings on cell and vacuole volume are pertinent for their use as the model unicellular eukaryotic organisms. W303 stood out for having a very high standard deviation for cells larger than the median in all tested conditions and its heterogeneity could be ascribed to the variability in vacuole volume. Our results show that the vacuole to cell volume ratio increases for W303 and KM, but not for CEN.PK and RT with the increase in cell volume indicating vacuole volume is relevant for the control of cell volume (S2B, S3B, S4B). This is important because cell volume is proportional to age and with aging the vacuole-linked autophagy becomes increasingly dysfunctional (56). The vacuolar dysfunction prohibits efficient catabolism of intravacuolar materials, induces enlargement of vacuole and, among others, can affect function of mitochondria and is implicated in the metabolic diseases (56). The reproducibility of lifespan extension results using yeast as model organism is a matter of concern (57). Our results suggest a careful consideration of the strain background while using yeasts in the aging research as different yeasts show the distinct vacuole to cell volume ratios that can potentially affect lifespan.

In conclusion, our study reports potassium and sodium salt tolerance responses in the non-conventional yeast vs. *S. cerevisiae* strains by describing their morphological and physiological features. As transport and efflux engineering is an important area of research in biotechnology our results will be relevant for further investigations concerning cationic transporters and stress tolerance in yeasts (58). Our open-source computational method for the optical image analysis will be a valuable community resource for cell and vacuole volume quantifications. A further benchmarking of the molecular mechanisms in the non-conventional yeasts compared with *S. cerevisiae* will be valuable for designing efficient MCFs with a broad substrate-product spectrum and a better tolerance of the industrially relevant cultivation conditions.

## Materials and Methods

### Strains and culture conditions

Yeasts used in the study included previously characterized strains of *Saccharomyces cerevisiae* CEN.PK113-7D (3), *S. cerevisiae* CJM 567 isogenic to W303 (59), *Kluyveromyces marxianus* CBS6556 (16) and *Rhodosporidium toruloides* CCT0783 (60). The genome and chromosomal these strains were as following: *S. cerevisiae* strains consisted of 12 megabase (Mb) genome distributed over 16 chromosomes, *K. marxianus* consisted of 11Mb genome spreaded over 8 chromosomes and *R. touloides* had 20Mb genome over 16 chromosomes (9, 10, 61, 62). All experiments were performed in a previously described minimal mineral medium containing 10 g of glucose, 5 g of (NH_4_)_2_SO_4_, 3 g of KH_2_PO_4_, and 0.5 g of MgSO_4_·7H_2_O per litre, in addition to 1 ml of trace elements solution and 1 ml of vitamin solution (63). The trace element solution contained, per litre (pH = 4), EDTA (sodium salt), 15.0g; ZnSO_4_·7H_2_O, 4.5g; MnCl_2_·2H_2_O, 0.84g; CoCl_2_·6H_2_O, 0.3g; CuSO_4_·5H_2_O, 0.3g; Na_2_MoO_4_·2H_2_O, 0.4g; CaCl_2_·2H_2_O, 4.5g; FeSO_4_·7H_2_O, 3.0g; H_3_BO_3_, 1.0g; and KI, 0.10g. The vitamin solution contained, per litre (pH = 6.5), biotin, 0.05g; p-amino benzoic acid, 0.2g; nicotinic acid, 1g; Ca-pantothenate, 1g; pyridoxine-HCl, 1g; thiamine-HCl, 1g; and myoinositol, 25g (63). The culture medium pH was adjusted to 6 with 1.9 ml per litre of 2M KOH. KOH was chosen because potassium ions were already being present at high enough concentration so that addition of KOH did not change the overall concentration significantly. Final K^+^ concentration in the minimal medium was 25mM. KCl and NaCl salts were used to achieve the required concentration of K^+^ and Na^+^. Physiology characterization experiments were conducted using 50mL flasks with vented caps having 20mL of culture medium. If necessary, for downstream processing, cells were washed with 0.9% saline solution. All strains were stored at −80°C in a mixture of 50% YPD (1% Yeast Extract, 1% peptone, 2% glucose) and 50% glycerol. For all the experiments, cells were pre-cultured in YPD overnight, washed and then grown in minimal medium with 1% glucose for 6h. Afterwards, cells were washed again and inoculated in a required medium to obtain initial OD of 0.1. Cultivation temperature was maintained at 30°C and mixing at 200 RPM allowed maintenance of aerobic culture environment.

### Cell density

Hitachi U-1800 spectrophotometer (Japan) and SARSTEDT polystyrene cuvettes were used for cell density measurements. The culture aliquots were diluted to keep OD_600nm_ in the range of 0.05–0.3 and the distilled water was used to set the reference.

### Metabolites measurements

0.25 mL of sample was collected for each data point of interest. In preparation for high-performance liquid chromatography, samples were centrifuged twice at 11,000g for 5 minutes and each time transferred to a new tube to remove the cells. Afterwards, samples were diluted 10 times to decrease the salt concentration and 60 μL of a diluted sample was used for measurements. As previously described the Aminex HPX-87H chromatography column (Bio-Rad) was used with the settings for elution of sugars and organic acids (temperature 45°C; flow rate 0.6 ml/min using mobile phase of 5 mM sulfuric acid) in a refractive index containing HPLC instrument (Prominence-I, LC-2030 C Plus, SHIMADZU) (63).

The carotene measurement was done according to previously published protocol (16). Samples (2 mL) were collected from stationary phase cultures of *R. toruloides* by centrifugation, washed twice in saline and resuspended in 1.0mL of acetone. The cells were lysed with acid-washed glass beads (400–650μm) in the FastPrep homogeniser (3x, 4m/s for 20 s) (MP Biomedicals, CA, USA). After centrifugation at 15,000g for 5min, the acetone solution containing carotene was collected and stored at 4°C until further quantification. The carotenes were quantified by using Acquity UPLC (Waters, MA, USA) equipped with a TUV detector (Waters, MA, USA) and C18 column (BEH130, 1.7 μm, 2.1 × 100 mm, Waters, MA, USA). A gradient of 80 to 100% acetone was used in the mobile phase with purified water at a 0.2mL/min flow rate. Carotene were detected was at 450nm and identified peaks were quantified with the β-carotene standard (Alfa Aesar, MA, USA).

### Microplate reader

Yeasts were pre-cultured as described above and each well in a 96-well plate was filled with 200μL of culture medium and inoculum provided a starting OD_600nm_ of 0.1. A BioTek Synergy MX microplate reader (United States) equipped with a shaker and temperature was set to 30°C and controlled by Gen5 ver. 2.04 software. The culture plate was scanned every 30min after 15s high speed shaking at an absorbance of 600nm. The absorbance data was exported to a MS Excel for further analysis.

### Growth rate

An exponential curve was fit to a region indicating an exponential growth of the culture and reproducible data were considered for calculations when an R^2^ of at least 0.95 was present. Slope of the curve was taken as a specific growth rate (h^−1^).

### Cell and vacuole volume quantification by image analysis

Yeasts were pre-cultured as described above and samples for optical imaging were collected during mid-exponential phase as determined by OD measurements. 1mL of culture was centrifuged at 3,000g for 1 minute to obtain a cells pellet. The pellet was resuspended in in 3μL of the same growth medium, which was transferred on to a glass slide. Nikon Eclipse Ci-L (Japan) microscope was used at Ph3 setting. All images were acquired by using Samsung Galaxy S7 Edge (South Korea). For image analysis a custom neural network-based analysis software pipeline was developed in MATLAB and used for segmenting cells in images. Cell and vacuole volume (fL) were estimated by constructing a 3D model based on a 2D mask where the third dimension was optimized for circularity. Median value was used as a representative measure of volume. Since the normal distribution was absent, a standard deviation represented in error bars was calculated independently for each half of volume distributions, which were split into two halves according to respective medians. For visualization, only 50% values of standard deviations were plotted for positive and negative directions from the median. Ratios of vacuole to cell volume were calculated by dividing the median of vacuole volume in each bin. If a bin had less than 30 cells, then its width was increased until it had 30 cells. The detailed open source code used for image analysis is available via GitHub and archived to Zenodo. In order to perform significance analysis, the Kruskal-Wallis test was used using MATALB.

### Cationic transporters analysis

NCBI nucleotide and protein BLAST were used to determine homology between genes and proteins (64). Following genome resources were used for obtaining DNA and proteins sequences for a comparative analysis: *S. cerevisiae* CEN.PK-113-7D (9), *S. cerevisiae* W303 (43), *K. marxianus* (12), *R. toruloides* (44).

## Acknowledgments

This project has received funding from the European Union’s Horizon 2020 research and innovation program under grant agreement No 668997, and the Estonian Research Council (grant PUT1488).

We thank Prof. Juana M. Gancedo and Prof. Carlos Gancedo (Instituto de Investigaciones Biomédicas ‘Alberto Sols’ CSIC-UAM, Madrid, Spain) for kindly providing isogenic W303 strain.

## Salt manuscript main figures

**Figure 1 [Reference] [S1A, S1B]**

A. Yeasts (morphology)
B. **1.** Specific growth rates **2.** Cell volume (fL) **3.** Vacuole volume (fL)
C. **1.** Cells heterogeneity (fL) **2.** Vacuoles heterogeneity (fL) **2.** Vacuole/Cell (fL/fL) **4.** Physiology

2. **Figure 2 [Potassium stress] [S1A, S2B]**

A. Relative growth rates
B. Physiology
C. Cellular and vacuolar heterogeneity distribution
3. **Figure 3 [Sodium stress] [S1A, S3A, S3B]**

A. Relative growth rates
B. Physiology
C. Cellular and vacuolar heterogeneity distribution
4. **Figure 4 [Sodium + Potassium stress] [S1A, S4A, S4B, S4C]**

A. Relative growth rates
B. Physiology
C. Cellular and vacuolar heterogeneity distribution
5. **Figure 5 [RT, Carotene]**

A. K^+^/Na^+^ transporters
B. Salt stress application

**Salt manuscript supplemental data:**

1. **S1A** – Cations in the Delft medium [Ref. – **Results - characterization section**]
2. **S1B** – Physiology table/graph [Ref. – **Figure 1** **to** **4**]
3. **S2A** – Morphology [Potassium] [Refers – **Figure 2**]
4. **S2B** – cells % vs fL, vacuoles % vs fL, vacuoles/cells (fL/fL) [Refers – **Figure 2**]
5. **S3A** – Morphology [Sodium] [Refers – **Figure 3**]
6. **S3B** - cells % vs fL, vacuoles % vs fL, vacuoles/cells (fL/fL) [Refers – **Figure 3**]
7. **S4A** – Cellular adaptation and density [Refers – Figure 4]
8. **S4B** – Morphology [Sodium + Potassium] [Refers – **Figure 4**]
9. **S4C** - cells % vs fL, vacuoles % vs fL, vacuoles/cells (fL/fL) [Refers – **Figure 4**]
10. **S5A-B** – Transporters – comparison [Refers – **Figure 5**, **Discussion**]

