## Supplementary material for "Characterization of potassium, sodium and their interactions effects in yeasts": S1A

**Table S1A: Cations (mM) concentration in the reference culture medium**

| Cation | Concentration (mM) |
|---|---|
| $K^+$ | 25.845 |
| $Mg^{2+}$ | 2.029 |
| $Na^+$ | 0.055 |
| $Ca^{2+}$ | 0.033 |
| $Zn^{2+}$ | 0.016 |
| $Fe^{2+}$ | 0.011 |
| $Mn^{2+}$ | 0.006 |
| $Co^{2+}$ | 0.001 |
| $Cu^{2+}$ | 0.001 |
