## Supplementary figures and images for "Characterization of potassium, sodium and their interactions effects in yeasts"

### S2A

S2A

Cellular morphology of yeasts in response to varying potassium concentrations

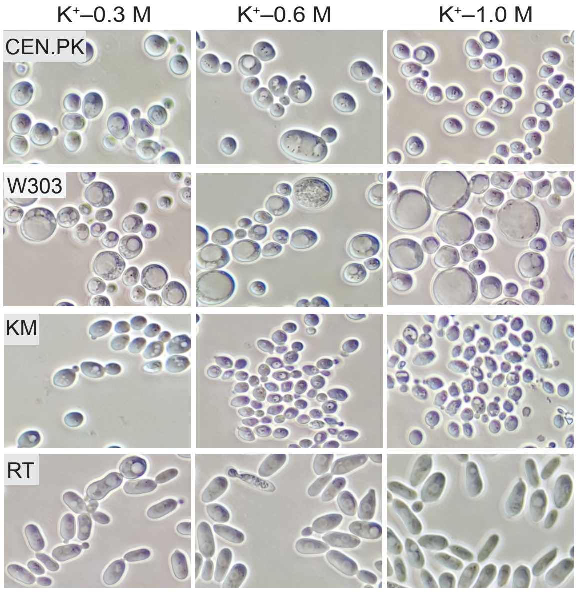

### S3A

# S3A

Cellular morphology of yeasts in response to varying sodium concentrations

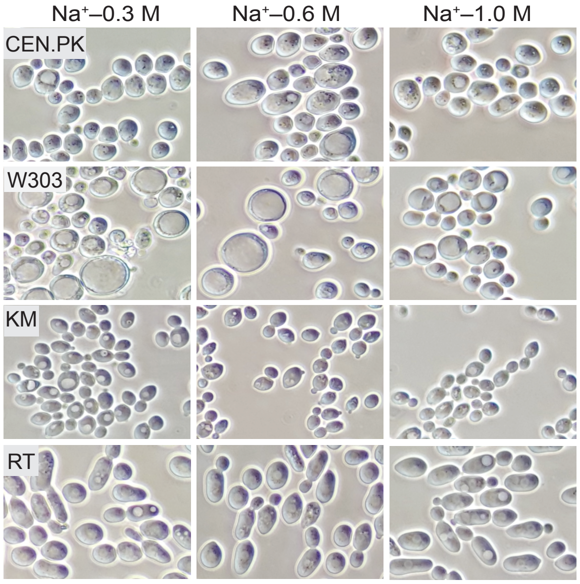

### S4B

S4B

Cellular morphology of yeasts in response to dual cationic stress

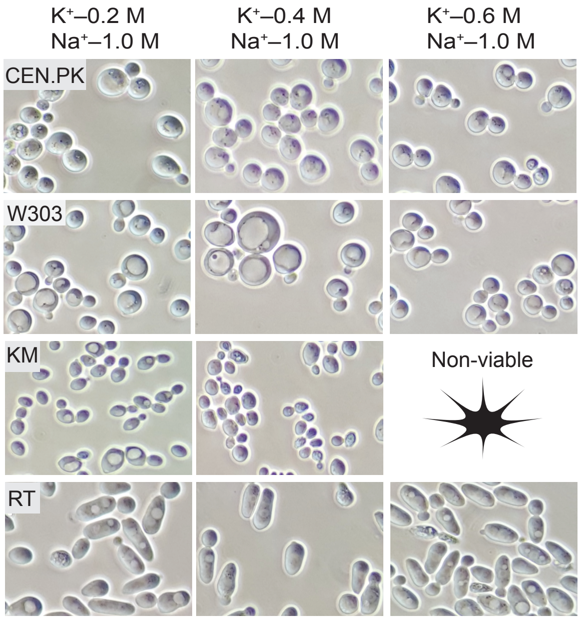
