## Supplementary material for "Characterization of potassium, sodium and their interactions effects in yeasts": S2B

Cellular and vacuolar volume distributions of yeasts in response to varying potassium concentrations. Cells and vacuoles numbers (n) used in the ratio calculations are the same as used for determining volume distributions for vacuoles.

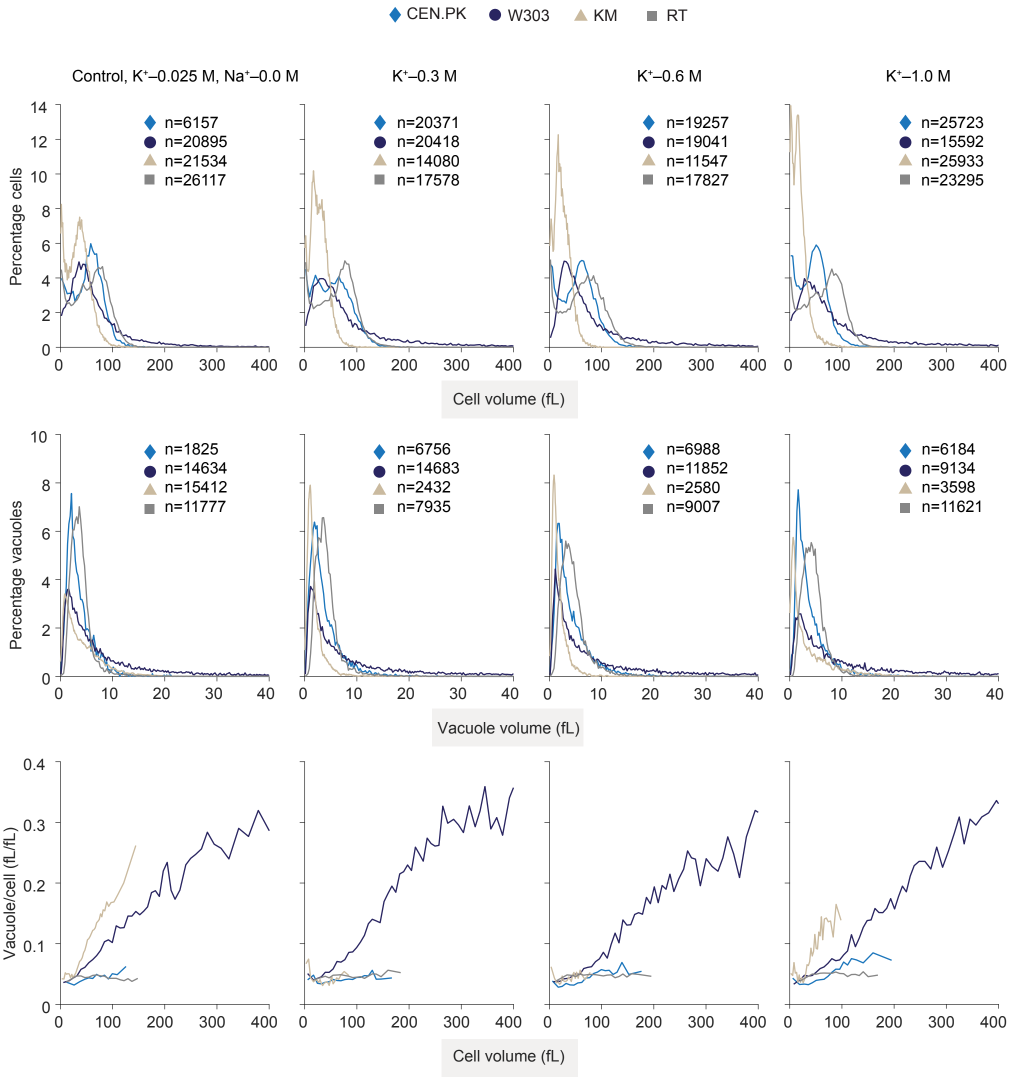
