## Supplementary material for "Characterization of potassium, sodium and their interactions effects in yeasts": S4A

Impact of salt stress on lag phase and
the final cells densities in response to dual cationic stress

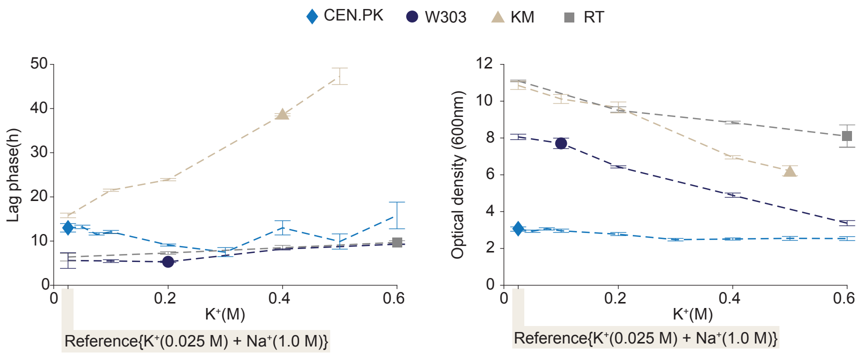


Impact of salt stress on lag phase

Final cells densities in shake-flask cultures
