## Supplementary material for "Characterization of potassium, sodium and their interactions effects in yeasts": S5A-B

**Table S5A. Comparison of sodium and potassium transporters in yeasts**

| Transporter Family | *S. cerevisiae* (W303) | *S. cerevisiae* (CEN.PK) | *K. marxianus* | *R. toruloides* |
|---|---|---|---|---|
| **PLASMA MEMBRANE** | | | | |
| **Influx** | | | | |
| Trk family ($K^+$) | 2 | 2 | 1 | 2 |
| Hak family ($K^+$) | - | - | 1 | 1 |
| Pho89 ($Na^+$) | 1 | 1 | 1 | 1 |
| Acu family ($K^+$/$Na^+$) | - | - | - | 2 |
| **Efflux** | | | | |
| Tok family ($K^+$) | 1 | 1 | 1 | 2? |
| Nha family ($Na^+$/$K^+$) | 1 | 1 | 1 | 2 |
| Ena family ($Na^+$/$K^+$) | 4 | 1 | 1-? | 1-? |
| **INTRACELLULAR** | | | | |
| **Influx** | | | | |
| Nhx family ($Na^+$/$K^+$) | 1 | 1 | 1 | 1 |
| Stv1 ($H^+$) | 1 | 1 | 1 | 1 |
| Vph1 ($H^+$) | 1 | 1 | 1 | 1 |
| Vnx1 ($Na^+$/$K^+$) | 1 | 1 | 1 | - |
| Vhc1 ($K^+$) | 1 | 1 | 1? | - |
| Kha1 ($K^+$) | 1 | 1 | 1 | 2 |
| Mdm38 ($K^+$) | 1 | 1 | 1 | - |
| Mrs7 ($K^+$) | 1 | 1 | 1 | 1 |

**Table S5B. A comparative analysis of monovalent cation transporters for yeasts used in the study. W303 is used as the reference strain. Number of differing nucleotides and amino acids are specified for CEN.PK strain.**

| | CEN.PK-113-7D | | *K. marxianus* | *R. toruloides* |
|---|---|---|---|---|
| | DNA identity | Protein identity | Protein identity | Protein identity |
| **Plasma membrane** | | | | |
| TRK1 | 99.27%<br>27 nucleotides | 98.87%<br>14 amino acids | 53.90% | 41.76% |
| TRK2 | 99.48%<br>13 nucleotides | 99.66%<br>3 amino acids | - | 42.70% |
| TOK1 | 100% | 100% | 37.28% | 34.65% &<br>23.59% |
| NHA1 | 100% | 100% | 50.60% | 40.50% &<br>37.61% |
| ENA | 90.80% with ENA2<br>310 nucleotides | 94.96% with ENA2<br>49 amino acids | 66.04%<br>with ENA2 | 24.52%<br>with ENA2 |
| PHO89 | 100% | 100% | 60.46% | 33.92% |
| **Intracellular** | | | | |
| Nhx1 | 100% | 100% | 64.81% | 50.98% |
| Stv1 | 99.5%<br>12 nucleotides | 99.78%<br>2 amino acids | 59.98% | 39.45% |
| Vph1 | 100% | 100% | 62.50% | 43.30% |
| Vnx1 | 100% | 100% | 55.44% | - |
| Vhc1 | 99.97%<br>1 nucleotide | 99.92%<br>1 amino acid | 57.52% (?) | - |
| Kha1 | 100% | 100% | 42.98% | 41.19% and<br>39.35% |
| Mdm38 | 99.94%<br>1 nucleotide | 99.83%<br>1 amino acid | 68.91% | - |
| Mrs7 | 100% | 100% | 69.30% | 43.17% |
